# Population responses of naïve roe deer to the recolonization of the French Vercors by wolves

**DOI:** 10.1101/560128

**Authors:** M Randon, C Bonenfant, J Michallet, T Chevrier, C Toïgo, J-M Gaillard, M Valeix

## Abstract

In a context of rapidly changing carnivore populations worldwide, it is crucial to understand the consequences of these changes for prey populations. The recolonization by wolves of the French Vercors mountain range and the long-term monitoring (2001-2017) of roe deer populations provided us a unique opportunity to assess both lethal and non-lethal effects of wolves on these populations. We compared roe deer population abundance and growth, fawn body mass, and browsing intensity in two contrasted areas: a central area (core of a wolf pack territory characterized by an intense use by wolves) and a peripheral area (used more occasionally). Both populations of roe deer strongly dropped after an extremely severe winter but the population of the central area facing with wolves was slower to recover (due to a much lower growth rate the following year) and remained at lower abundance levels for 5 years. Fawn body mass was lower in the central area during that period and was not influenced by weather conditions or population abundance. The browsing index in the forests in presence of wolves decreased for a longer period, suggesting that possible habitat shifts have occurred. Altogether, the effects of wolves on the roe deer population in the central area occurred mainly during a 5-year period following the establishment of wolves, with effects at the population level in the first years only through the interplay between wolf predation (before wolves started preying on red deer), harsh winter conditions and naïveté of prey to this recolonizing predator.

## Introduction

The main drivers of population dynamics of large herbivores have been studied into much details over the last decades (e.g. Coulson et al. 2001; Gaillard et al. 2013 for case studies). The consequences of density, weather, habitat quality or hunting on age-specific survival and reproduction are well documented in many species of large herbivores, with increasing empirical evidence of interactions among those limiting factors (Hone and Clutton-Brock 2007; Bonenfant et al. 2009). Predation is clearly a major driver of evolution and population dynamics of prey (Volterra 1931; Reznick et al. 2004). Understanding and measuring the consequences of predation on the population dynamics of large herbivores is, however, much more complex than for most other environmental variables. Consequently, important ecological questions such as whether large herbivores are undergoing a bottom-up or to-down limitation are still debated (Hopcraft et al. 2010; Laundré et al. 2014).

By killing prey and increasing mortality, predators are strongly expected to limit the population growth rate of their prey but there are several arguments suggesting that prey populations can support strong predation pressure. If mortality from predation is compensatory because of density-dependence, population dynamics of prey may remain little affected by predation until attack rates become really high and mortality from predation becomes additive to other sources of mortality (Errington 1946). Similarly, the difference in spatial scale between the ranging behaviour of large carnivores and herbivores leads to differences in densities of several orders of magnitude between predators and prey (Skogland 1991). Consequently, large predators may have limited consequences for population growth rate of prey and particularly so if predators are generalists and can switch between different prey species (Murdoch 1969) or if they select juvenile or senescent individuals because in large herbivores, the population growth rate is most sensitive to variation in the survival of prime-aged adults (Gaillard et al. 2000). However, highly specialized predator species or individuals can clearly reduce population growth rate and the abundance of large herbivores (Festa-Bianchet et al. 2006; Bourbeau-Lémieux et al. 2011). For instance, roe deer *Capreolus capreolus* dynamics is markedly affected by lynx *Lynx lynx* predation (Heurich et al. 2012; Andrén and Liberg 2015) particularly so in winter when snow layer is thick, which greatly limits roe deer mobility (Heurich et al. 2012).

In a context of rapidly changing abundance and distribution of mammalian apex carnivore populations worldwide (Chapron et al. 2014; Ripple et al. 2014), it is important to understand the consequences of these changes for prey populations and ultimately for ecosystem functioning. Even though studies on these consequences have accumulated over the past decades, most of our current knowledge comes from studies from North American National Parks (and particularly from the grey wolf *Canis lupus* and elk *Cervus canadensis* of the Greater Yellowstone Ecosystem) as pointed out by Kuijper et al. (2016). There is thus a need for studies from different contexts, particularly in Europe where large carnivores live in or are recolonizing anthropogenic landscapes (Chapron et al. 2014). Further, whether prey have continuously co-evolved with their predator or have evolved in a predator-free environment for several generations due to predator extirpation from some ecosystems may ultimately influence the extent to which prey are vulnerable to predators (Byers 1997; Berger et al. 2001). Indeed, naïve prey may fail to recognise the cues of a novel predator (but see Chamaillé-Jammes et al. 2014) or may fail to respond appropriately and effectively to the risk of predation by this predator due to the lack of experience (Banks and Dickman 2007; Carthey and Banks 2014). For instance, along brown bear *Ursus arctos* recolonization fronts, brown bears killed adult moose *Alces alces* at disproportionately high rates compared to sites where brown bears have always been present (Berger et al. 2001). However, very little is known on how naïve prey respond to recolonizing predators and how quickly they become effective at efficiently escaping these predators.

In 1992, wolves crossed the Italian border to recolonize eastern France from where the predator had been missing for ca. 100 years (Valière et al. 2003). In this work, we took advantage of the long-term monitoring (17 years) of roe deer populations in the west Vercors mountain range covering contrasting areas in terms on wolf occupancy and abundance to assess the occurrence and relative impact of lethal and non-lethal effects of wolves on these roe deer populations, and the duration of the naïve period in these roe deer populations. If predation by wolves and the associated predation risk affect roe deer, we expect (1) a decrease in the roe deer population abundance and growth rate, (2) a decrease in roe deer fawn body mass, and (3) a decrease in the herbivore pressure on the vegetation, following the return of wolves.

## Materials and Methods

### Study area

The study was carried out between 2001 and 2017 over a study area of 30 776 ha encompassing six neighbouring counties (Bouvante (8 431 ha), La Chapelle-en-Vercors (4 562 ha), Vassieux-en-Vercors (4 798 ha), Saint-Julien-en-Vercors (1 867 ha), Saint-Martin-en-Vercors (2 700 ha), Saint-Agnan-en-Vercors (8418 ha)) in the French department of Drôme, in the west Vercors mountain range (Fig. 1). The west Vercors mountain range is characterized by an Alpine climate (identical to the Northern Alps) with a forest dominated by beech *Fagus sylvatica* and silver fir *Albies alba*. The large herbivore community is composed of roe deer, red deer *Cervus elaphus*, chamois *Rupicapra rupicapra*, mouflon *Ovis gmelini*, and wild boar *Sus scrofa*. The six counties are used by people for agriculture, livestock breeding, forestry, hunting, and outdoor recreational activities.

**Figure 1:**
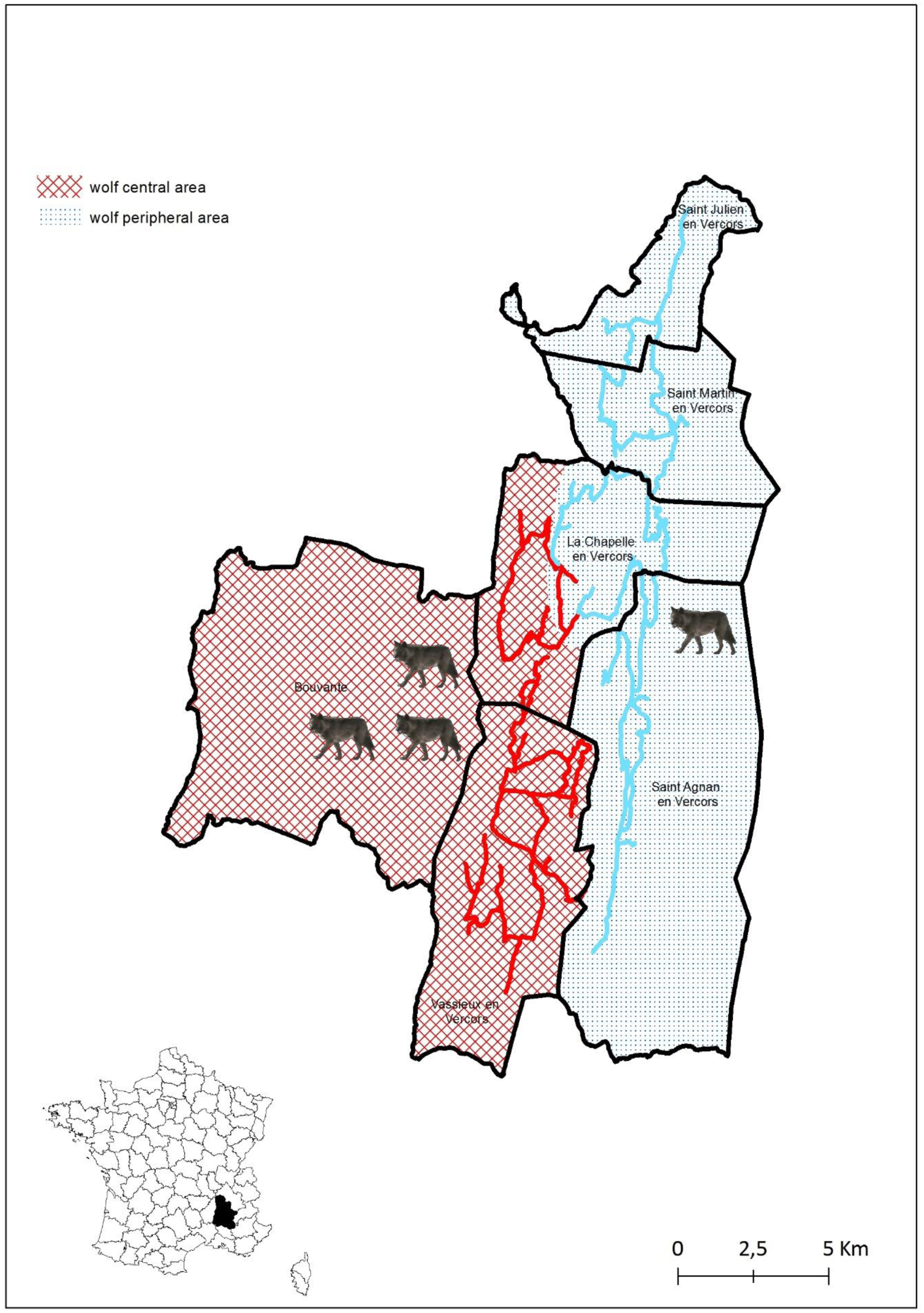
Map of the study area in the French department of Drôme with the location of the six study counties (the small map of France in the left bottom corner shows the location of the Drôme department in France). The red hatched area represents the central area of the west Vercors wolf pack territory, while the dotted blue area represents the peripheral area. Coloured lines show the transects for the monitoring of roe deer population abundance.

### History of wolf presence in the west Vercors mountain range

Wolves were extirpated from the Drôme department in 1901 (Faton and Ladreyt 1982). In 1998, the first field evidence from prey carcasses, tracks and faeces suggesting the return of wolves from the Italian Alps to the west Vercors mountain range were found (Valière et al. 2003). The ONCFS (French National Hunting and Wildlife Agency) network “Grands Prédateurs” later confirmed the permanent occupancy (3 individuals identified based on DNA analyses) and reproduction of wolves in the west Vercors mountain range in 2003/2004 (ONCFS 2006). At this early stage of the recolonization, only lone wolves or single pairs were observed. Since 2007/2008, wolves form packs of a minimum of five individuals. In this study, we contrasted two main study areas based on the intensity of use by wolves. The central area is the core of the west Vercors wolf pack territory (central area hereafter), and encompasses the counties of Bouvante, Vassieux-en-Vercors and the western sector of La Chapelle-en-Vercors (Fig. 1). The central area is characterized by an intensive use of the area by wolves where sightings of wolves, wolf tracks, and wild prey carcasses are frequently reported. In contrast, the peripheral area is used by wolves more occasionally, and encompasses the eastern sector of La Chapelle-en-Vercors, Saint-Julien-en-Vercors, Saint-Martin-en-Vercors, and Saint-Agnan-en-Vercors (Fig. 1).

### Weather data

We obtained weather data (daily rainfall and mean daily temperature) from Météo France for the weather station La Chapelle-en-Vercors, which is representative of the west Vercors mountain range. We calculated the Gaussen index (i.e. the amount of precipitation minus twice the mean temperature) to measure the water deficit of plants in spring (April-June) and summer (July-August) (e.g. Gaillard et al. 1997; Garel et al. 2004) to which roe deer are particularly susceptible (Pettorelli et al. 2005). The Gaussen index is a proxy of the balance between rainfall and evapo-transpiration of plants (Gaussen and Bagnouls 1953). High values of the Gaussen index mean positive water balance, higher plant growth, and hence better foraging conditions for large herbivores, and conversely (Toïgo et al. 2006). Winter can be very long in the west Vercors mountain range so this season was defined from October to March. To characterize winter conditions, we collected information on snow-fall, snow depth, and number of days with snow cover from the local skiing resorts in Bouvante. Because of the strong correlation existing among winter variables, we performed a principal component analysis (PCA) on these standardized variables. The first principal component (PC1) accounted for 62% of the overall variance, so we used the point projections on PC1 as a winter harshness index. Low values of this index were associated with severe winters and hence more difficult conditions for roe deer as harsh winters are generally associated with a lower survival of fawns (Gaillard et al. 1993) and costly movements for large herbivores (Parker et al. 1984), which, in turn, could increase predation rates (Mech et al. 2001).

### Roe deer population abundance

We monitored the abundance of roe deer populations in 5 of the 6 study counties because Bouvante could not be monitored due to the deep snow cover that made most roads in this county inaccessible at the time of surveys in all years. We drove along 3 transects located in the central area (1 transect in the western sector of La Chapelle-en-Vercors and 2 transects in Vassieux-en-Vercors) and 3 transects in the peripheral area (1 transect in Saint-Agnan-en-Vercors, 1 transect across St-Julien-en-Vercors and St-Martin-en-Vercors, and 1 transect in the eastern sector of La Chapelle-en-Vercors – see Fig. 1). We carried out counts at night with a powerful spotlight reflecting animals’ eyes. We counted roe deer after winter, in March-April, when vegetation flush has not started yet, along roads known to be practicable at that time of the year. We drove transects at low speed (10-15 km/h) with one driver, two observers who spotted and identified all animals seen, and one person who recorded the observations. We repeated counts twice a year between 2001 and 2004, three times a year between 2005 and 2012, and four times a year since 2013. For the central and peripheral areas, we obtained an abundance index of roe deer population (AI) by calculating the mean number of roe deer seen per kilometre (see Pellerin et al. 2017 for a similar approach applied to diurnal car counts). Finally, we derived the annual growth rate (*r*_t_) of the yearly abundance indices as follows: *r*_t_ = ln(AI_t+1_/AI_t_). We assumed that this growth rate provided a reliable proxy of roe deer population growth from year *t* to year *t+1*.

### Roe deer fawn body mass

Twenty local hunting associations (which encompass 500 hunters) contributed to this study and were equipped with a digital scale with an accuracy of 100 grams to weigh hunted roe deer. Between 2002 and 2007, hunters measured the full body mass of harvested roe deer, but have switched to dressed body mass (i.e. guts, liver, heart and lungs removed) since 2007. Between 2007 and 2009, 43 local hunting associations in the whole Drôme department were asked to measure both full and dressed body masses. From a sample of 170 roe deer with the two measurements, we checked that a close relationship existed between dressed and full body masses (dressed body mass = (0.837 x full body mass) - 1.054; R² = 0.92) and used this relationship to estimate dressed body mass of roe deer harvested during 2002-2007. We used dressed body mass in all subsequent analyses. Because roe deer are income breeders with limited fat reserves (Andersen et al. 2000), and variation in adult body mass is mainly caused by early-life conditions (Pettorelli et al. 2002), we analyzed body mass of roe deer fawns (individuals < 1 year when shot). We excluded body mass data from la Chapelle-en-Vercors because the exact locations of where animals were shot were not recorded, which prevented us from assigning the hunted roe deer of this county to the central vs. peripheral area.

### Herbivore pressure on the woody vegetation

Because changes in browsing pressure correlate with changes in the abundance of populations of large herbivores (Morellet et al. 2001; Chevrier et al. 2012), we monitored the browsing pressure in the forest habitats of the central area of the wolf pack territory (Bouvante, Vassieux-en-Vercors, and the western sector of La Chapelle-en-Vercors) from 2001 to 2014. Unfortunately, such monitoring did not take place in the peripheral area. One limit of such index is that it encompasses the browsing pressure from all herbivore species. In the study system, this index encompasses the browsing pressure from both roe deer and red deer. For this monitoring, we focused on the four main woody plant species of the west Vercors mountain range (beech, silver fir, Norway spruce *Picea abies*, and sycamore *Acer pseudoplatanus*). Every year in April-May, between snow melt and spring vegetation flush, we monitored 86 quadrats (1 m²) distributed in the central area. In each quadrat, we recorded whether one of these four species was present and whether these plants had been browsed in the past growing season. Following Morellet et al. (2001), the browsing index was defined as *B* = (n_c_ + 1)/(n_p_ + 2) where n_p_ is the number of plots where at least one of the monitored species was present, and n_c_ is the number of plots with at least one species consumed.

## Analyses

### Roe deer abundance

We analysed variation in roe deer abundance (assessed using the AI) with Generalised Linear Models (GLMs) setting a logarithmic link function and a negative binomial distribution. We opted for a negative binomial distribution because the model with a Poisson distribution did not fit the data well (goodness-of-fit test: χ² = 1 451.71, df = 287, P < 0.001) resulting from over-dispersed count data (ver Hoef and Boveng 2007). Even if we did our best not to change the road count protocol, transect length did vary among years and across transects. Including an offset variable (log-transformed number of kilometres) accounted for this heterogeneity in the data collection. We included a categorical variable ‘year’ with 16 levels to test for temporal variation in roe deer abundance. We investigated whether the temporal dynamics of roe deer abundance differed between the central and peripheral areas by testing the first-order interaction between the effects of ‘year’ and ‘wolf area’ (a 2-level categorical variable: “central area” and “peripheral area”). Finally, we fitted a model where environmental covariates accounted for temporal variation in roe deer abundance and tested the prediction that predation and winter conditions lead to decreasing population abundance with the interaction term between the effects of the ‘winter harshness index’ and ‘wolf area’. For the population growth rate (*r* assessed from AI), we investigated the time variation using Gaussian linear models; ‘year’ being entered as a categorical variable. We also fitted a model with first-order interaction term between the effects of ‘year’ and ‘wolf area’. For both AI and population growth rate *r* assessed from AI, we assessed the statistical significance of all variables with likelihood-ratio-tests (LRT).

### Fawn body mass

We analysed fawn body mass of roe deer using Gaussian linear models. All models, consistently included both sex (a 2-level categorical variable) and date of harvest (the number of days elapsed since June 1^st^ of the year of birth) as explanatory variables to account for fawn body growth over the hunting season. We tested for temporal variation in average body mass of fawns by including ‘year’ as a categorical variable. The interaction term between the effects of ‘year’ and ‘wolf area’ was also included to test for possible difference in time variation in fawn body mass between the central and peripheral wolf areas. We then quantified and tested for the effects of spring and summer Gaussen index at year *t*, and of winter harshness for the winter season covering years *t* and *t+1* on fawn body mass from the hunting season covering years *t* and *t+1* by replacing year with the corresponding weather index in the model, one at a time. Again, we included the interaction term between the effects of ‘wolf area’ and weather indices. For our model selection, we sequentially removed non-statistically significant variables starting from the most complex model. We tested for the effect of sex, date of harvest and year using LRT. For the effect of environmental covariates, we tested their significance using an analysis of deviance (ANODEV) (Skalski 1996; Grosbois et al. 2008).

### Browsing index

The browsing index can reasonably be approximated by a proportion of consumed plants by herbivores. We hence modelled the browsing index using GLMs, with a logistic link function and a binomial distribution to constrain the response variable within the 0–1 interval. Our model included a random effect of the plot identity because of repeated measurements at the same place over years. We found no evidence for spatial correlation among the plots, so we did not specifically model plot location. We then added a year effect (a 14-level categorical variable) to test for an effect of time on the average browsing rate. Again, we assessed the statistical significance of time variation in the browsing index using LRT tests.

We performed all analyses with the statistical software R 3.4 (R Core Team 2018) extended with the *MASS* package (Venables and Ripley 2002). We set the significance level to α = 0.05 and reported estimates as mean ± 95% confidence interval unless otherwise stated.

## Results

Weather variables (i.e. winter harshness index, spring and summer Gaussen indices) during the study period are provided in Appendix 1. Winter 2004/2005 was the harshest of the time series, with a record of snow-fall (total snow-fall = 498 mm, max snow depth = 140 mm) and snow duration (number of days with snow cover = 110).

### Roe deer population abundance and growth rate

The negative binomial model fitted the data satisfactorily (goodness-of-fit test: χ² = 292.55, df = 287, P = 0.40). The AI of the roe deer population varied a lot between 2001 and 2017 (χ² = 97.60, df = 16, P < 0.0001), with different patterns between the central and peripheral areas (interaction between ‘year’ and ‘wolf area’: χ² = 41.39, df = 16, P < 0.0001; Fig. 2A). Roe deer AIs were positively associated between the central and peripheral areas except during the 2005-2010 period. In 2005, roe deer AI decreased dramatically in both areas, coinciding with the most severe winter of the study period (2004-2005). Between 2005 and 2010, roe deer AI remained low in the central area while it increased in the peripheral area (Fig. 2A). Since 2011, the annual variation in roe deer AI was synchronous in the two areas, as for the period 2001-2005 (Fig. 2A). Winter harshness did not account for temporal variation observed in roe deer AI (χ² = 0.10, df = 16, P = 0.99).

**Figure 2:**
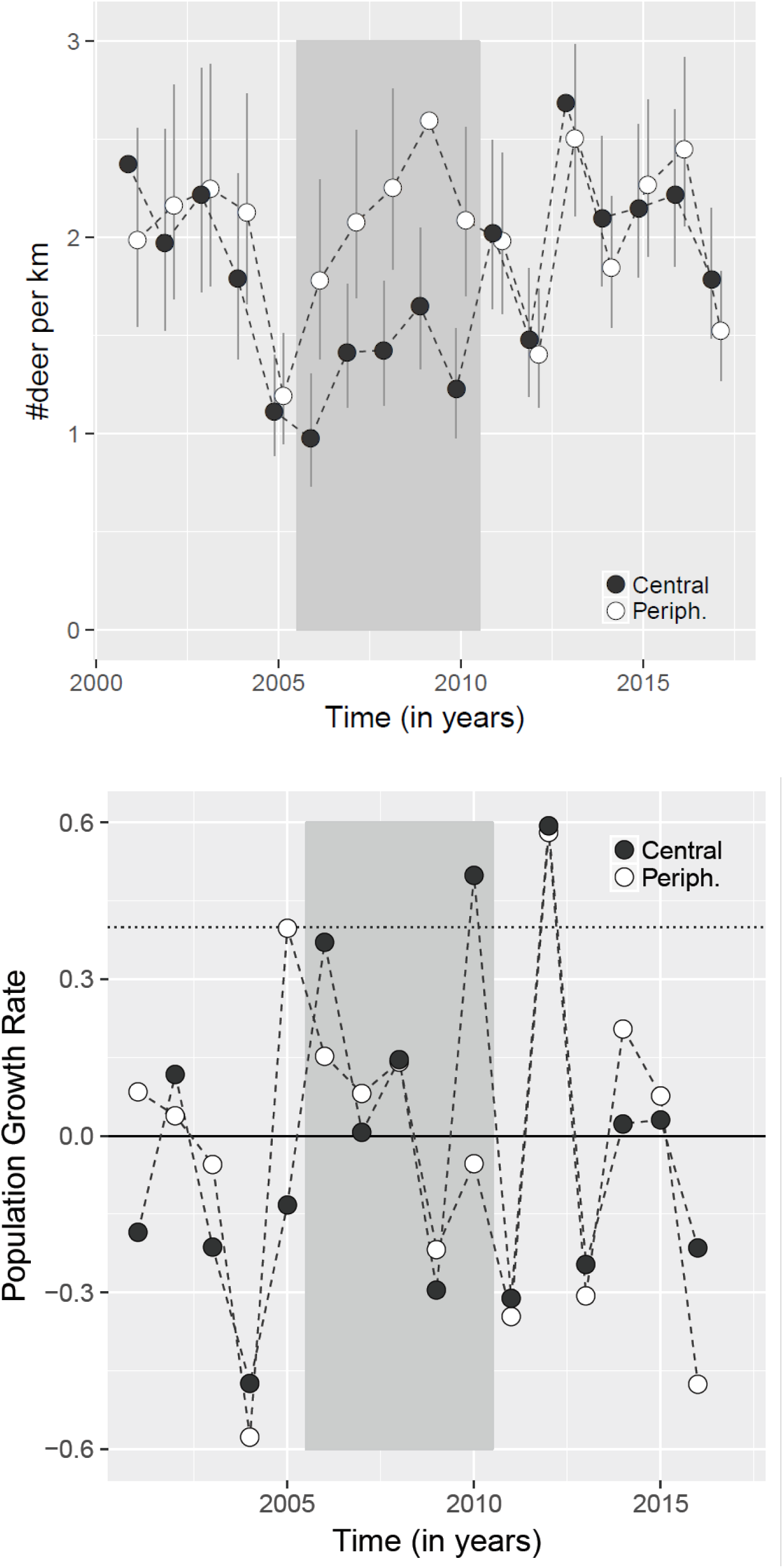
**A)** Changes in the abundance kilometric index of the roe deer populations in the central and peripheral areas of a wolf pack territory in the French west Vercors mountain range for the period 2001-2017. Bars represent the 95% confidence interval. **B)** Changes in the population growth rates of the roe deer populations in the central and peripheral areas of a wolf pack territory in the French west Vercors mountain range for the period 2001-2017. In A and B, the grey area shows the period for which the abundance kilometric indices are different between the central and peripheral areas.

All annual growth rates of AI were biologically realistic (i.e. < 0.4) except for two years (Fig. 2B), which could be explained either by methodological biases (e.g. exceptional local conditions during the counts) or by immigration. Figure 2B shows that the difference in roe deer AI between the two areas for the period 2005-2010 resulted from the much lower growth rate in the central wolf area between 2005 and 2006 compared to the peripheral area. After 2006, the annual growth rates were rather similar in the two study areas (Fig. 2B). Despite the lowest growth rate occurred between 2004 and 2005 henceforth including the harshest winter, we could not detect any effect of any weather variable on annual growth rates over the study period (spring Gaussen index: β = −0.00019 ± 0.00057, t = −0.34, P = 0.74; summer Gaussen index: β = 0.00036 ± 0.00061, t = 0.59, P = 0.56; winter harshness index: β = 0.00018 ± 0.00038, t = 0.474, P = 0.64).

### Roe deer fawn body mass

Overall, we collected dressed body mass measurements for *n* = 428 roe deer fawns in the study area from 2002 to 2016, both in the central (Vassieux-en-Vercors, Bouvante; n = 243) and the peripheral (Saint-Julien-en-Vercors, Saint-Martin-en-Vercors, Saint-Agnan-en-Vercors; n = 185) areas. The mean difference in fawn body mass between sexes was 372 ± 230 g (males heavier, as expected; Douhard et al. 2017), and fawns gained on average 250 ± 90.3 g per month over the hunting season from September to the following March. Mean body mass of fawns varied among years (*F* = 1.10, df = (14, 410), P < 0.01) but not differently between the central and peripheral wolf areas (interaction term between ‘year’ and ‘wolf area’: *F* = 1.10, df = (12, 398), P = 0.35). Fawn body mass was on average lower in the central than in the peripheral wolf area (0.700 ± 0.253 kg, *t* = 2.75, P < 0.01), consistently between 2006 and 2011 but no clear patterns were detected before and after that period (Fig. 3). The differences in mean fawn body mass among years and between areas were, however, rather low (≤ 1 kg). Of the four environmental covariates (i.e. the 3 weather indices and the population abundance index), none accounted for annual variation in fawn body mass (Table 1).

**Figure 3:**
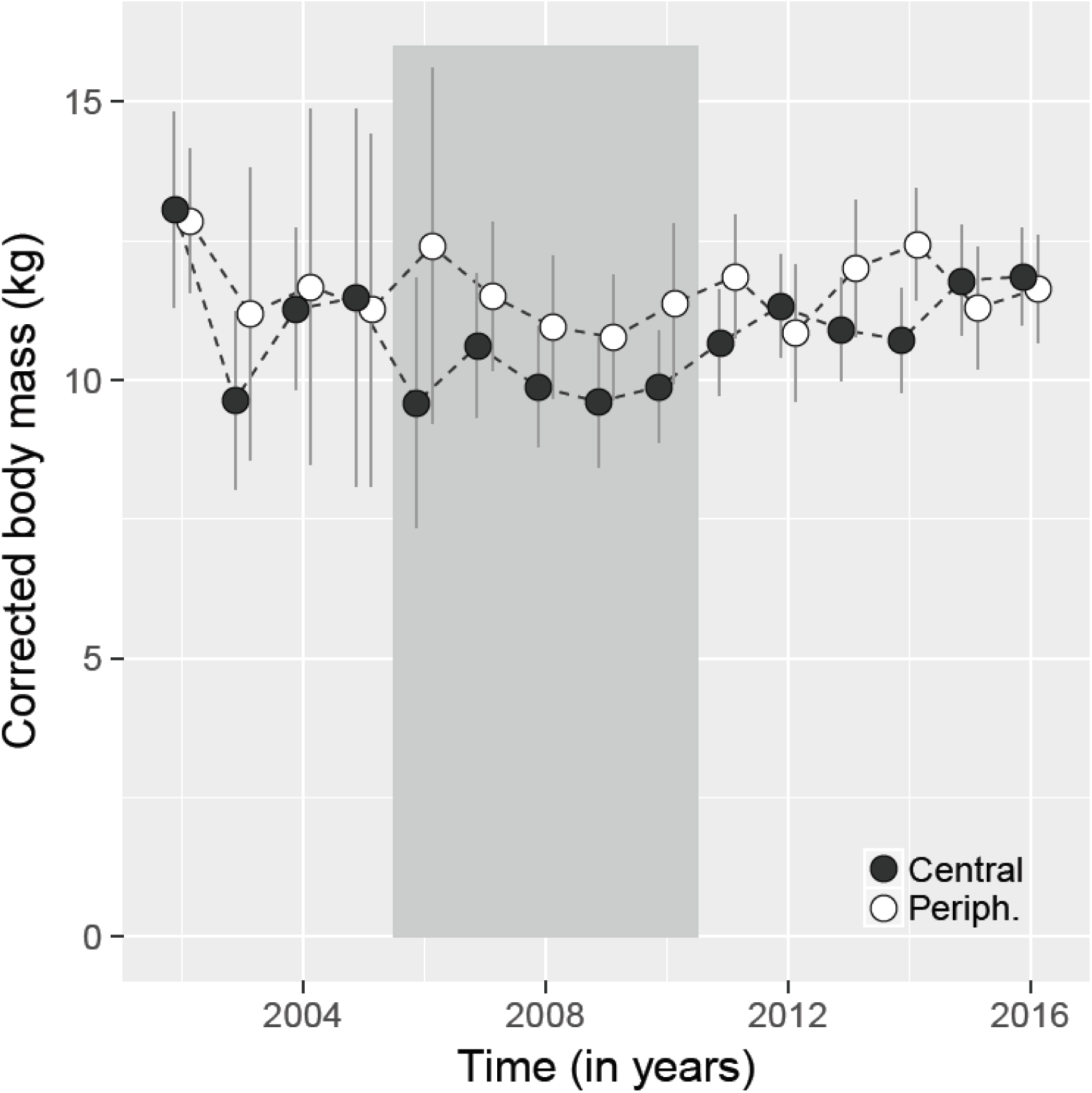
Changes in the roe deer fawn body mass (corrected dressed body mass – see text for details) in the central and peripheral areas of a wolf pack territory in the French west Vercors mountain range. Bars represent the 95% confidence interval. The grey area shows the period for which the body masses are different between the central and peripheral areas.

**Table 1:** Results from the analysis of deviance of the effect of environmental covariates on roe deer fawn body mass in the west Vercors mountain range (both the central and peripheral areas of the wolf pack territory). Data from dressed body mass measurements for *n* = 428 roe deer fawns harvested between 2002 and 2016.

| Environmental variable | Estimate | R<sup>2</sup> | ANODEV |
| --- | --- | --- | --- |
| Winter harshness index | -0.010 (0.118) | 0.00 | F = 0.004, df = (1, 13), P = 0.95 |
| Spring Gaussen index | 0.089 (0.113) | 0.07 | F = 1.17, df = (1, 14), P = 0.28 |
| Summer Gaussen index | 0.136 (0.113) | 0.06 | F = 0.84, df = (1, 14), P = 0.37 |
| Population abundance index | 0.215 (0.118) | 0.13 | F = 2.06, df = (1, 14), P = 0.17 |

### Herbivore pressure on the vegetation

The mean browsing index varied among years, between 0.10 in 2008 and 0.54 in 2001 (*F* = 22.27, df = (13, 5046), P < 0.01; Fig. 4). In the central wolf area, the browsing indices markedly decreased in the Vercors forests between 2001 and 2008, and continuously (but moderately) increased afterwards (Fig. 4).

**Figure 4:**
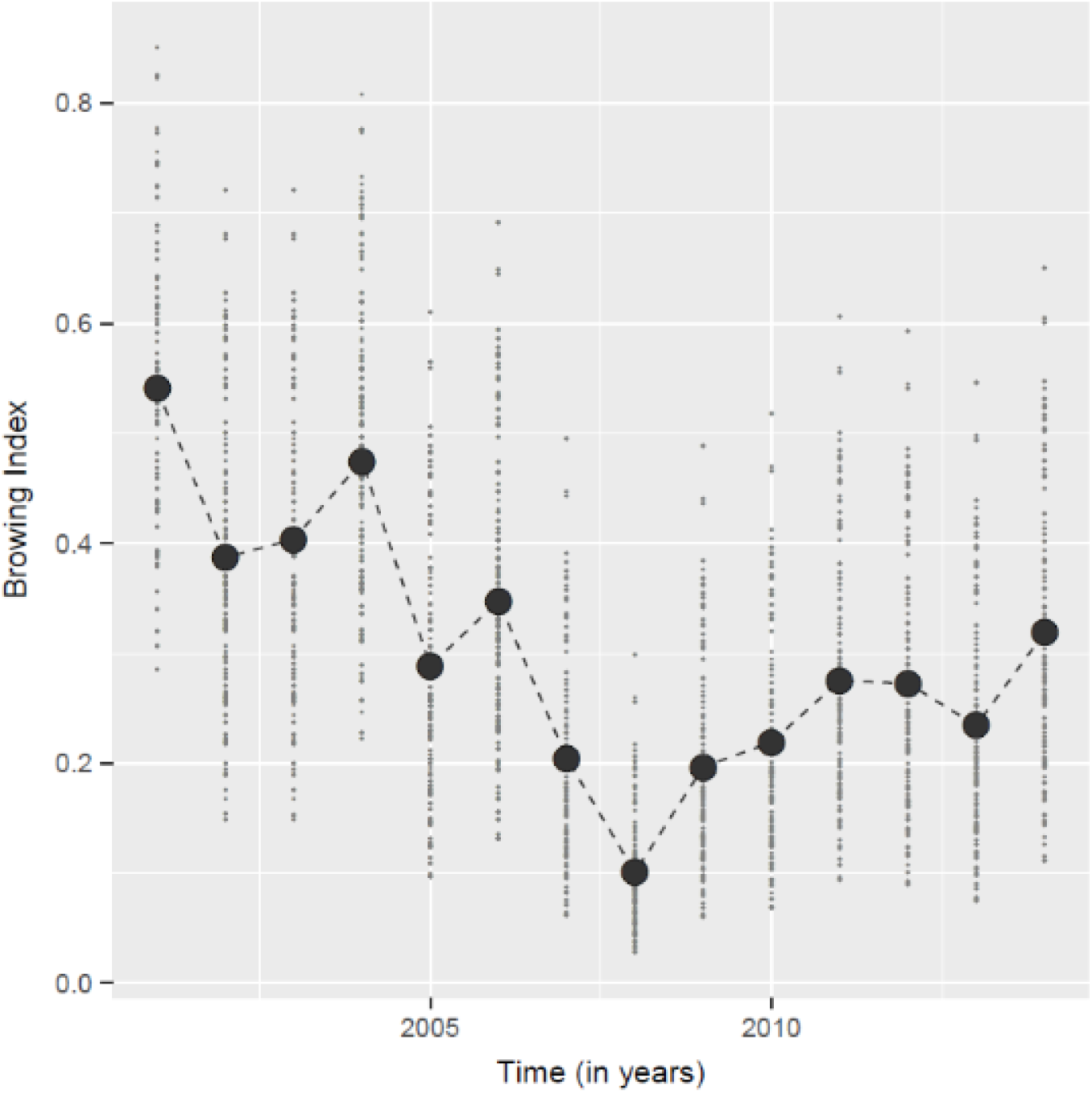
Changes in the browsing index (calculated for the four main woody plant species, i.e. beech, fir, Norway spruce, and sycamore) in the central area of a wolf pack territory in the French west Vercors mountain range.

## Discussion

Over the 17 years of our study, the roe deer population abundance patterns, as measured by the AI, were similar in the central and peripheral areas but during the 5-year period between 2006 and 2010. During that period, roe deer abundance was lower in the central area, which corresponds to the core of a wolf pack territory and is characterized by an intense use of the area by wolves. Populations of roe deer strongly dropped between 2004 and 2005 in both the central and the peripheral areas most likely because of the extreme severity of the winter 2004-2005, which was the harshest winter throughout the 17-year study period. This is consistent with several previous studies that showed that severe winters decrease survival of young and old individuals in populations of large herbivores (e.g. Saether 1997; Gaillard et al. 2000 for reviews). Wolf predation is expected to increase with snow depth. For instance, on the Isle Royale, wolves hunted in larger packs and tripled the number of moose they killed per day in the snowiest years (Post et al. 1999). Likewise, the relative importance of wolf predation on white-tailed deer *Odocoileus virginianus* mortality increased with winter severity in Minnesota (DelGiudice et al. 2002). Such higher susceptibility of ungulates to wolf predation during severe winters is associated with costlier and less efficient movements if ungulates in deep snow (Parker et al. 1984).

The steep decrease of roe deer AI occurred in both the central and peripheral areas, but the roe deer population was slower to recover in the central area. Something different occurred between the two study areas during the period 2006-2010. As no change in forest management occurred (Randon, pers. obs.), we can discard big change in resource availability to account for such a difference. Likewise, possible competition with red deer (Richard et al. 2010) cannot be involved because no detectable change in the red deer population abundance was detected during that period (unpublished data; supported by the lack of increase of the browsing index). The two wolf areas being very close geographically, differences in local weather conditions can also be excluded. No disease outbreak was reported over the study period (Randon, pers. obs.). The yearly variation in roe deer harvest bags was also similar in the two areas (Appendix 2). Predation by wolves is thus a likely factor to explain the difference we observe in population dynamics between the central and peripheral areas. This difference in population dynamics between 2006 and 2010 is due to a marked difference in the growth rate reported in the AI between 2005 and 2006 only. Because growth rates were not generally lower in the central area in the following years, this means that the mortality caused by wolf predation on roe deer in the central area influenced roe deer populations for 2 years only (2005 in interaction with the extreme winter severity and 2006 possibly still benefiting from the naïveté of roe deer) and after may not be large enough to be captured in the abundance index.

The sample sizes of roe deer fawn body mass were rather low but allowed us to depict a period (2006-2011) when body mass was lower in the central area than in the peripheral area. The positive relationship between roe deer fawn body mass and roe deer AI we reported is opposite to what was expected in presence of density-dependence (Bonenfant et al. 2009). Indeed, fawn body masses were lower in 2006-2010 when roe deer AI was low abundance in the central area. Such positive relationship has already been demonstrated in a study whereby bighorn sheep *Ovis canadensis* lambs suffered mortality through reduced growth during years of high predation by cougars *Puma concolor*, contributing a third of the total impact of predation on lamb survival (Bourbeau-Lémieux et al. 2011); a study that illustrated a case of non-consumptive effects of predation on a prey population. While our results may suggest such a mechanism, the alternative of a delayed effect of the extremely rigorous winter 2004/2005 that led several consecutive cohorts to be light, and hence prevented any relationship between average fawn body mass and population AI cannot be discarded.

The changes of the browsing index in the central area indicates a decrease of the browsing pressure on woody plants in forest habitats of the west Vercors mountain range. This decreased deer pressure on vegetation may result from either a decline in deer abundance (which does not seem to be the case when considering temporal variation in AI) or a change in foraging behaviour with an increased use of suboptimal habitats (as shown in other systems, e.g. Creel et al. 2005; Valeix et al. 2009). This may be the case here but future studies involving detailed GPS monitoring of individual roe deer are needed to investigate whether they alter their habitat use and selection as a response to predation risk by wolves. The risk of predation alone can affect prey individual performance through stress-mediated and food-mediated costs (Creel 2018; MacLeod et al. 2018). A study on wolves in the Greater Yellowstone Ecosystem, where wolves are wide-ranging and active predators, revealed that the behavioural responses of elk were not strong or frequent enough to lead to major changes in individual performance and, hence, on population abundance (Middleton et al. 2013).

This may be the case in the present study because behavioural adjustments of roe deer to the risk of predation by wolves are likely to carry costs at the individual level. However, it is noteworthy that the difference in fawn body mass was low (~1kg) compared to differences previously reported in roe deer in response to changes in density (about 2 kg in response to spatial variation in resources, Pettorelli et al. 2003; > 3kg in response to population density, Douhard et al. 2013)) but these ones do not translate into detectable changes in growth rates of AI, which are similar in the two contrasting study areas.

Roe deer populations in the central and peripheral areas had similar patterns of temporal variation of AI, growth rates and fawn body mass after 2011. This suggests that the effects of wolves on the roe deer population in the central area occurred mainly during a five-year period following the establishment of the pack, with effects at the population level in the first years only in interaction with the harshest winter in 2004/2005. The little difference we reported between the central and peripheral areas after 2011 may be explained by (i) a learning process to recognize wolf cues allowing roe deer to escape from wolf predation (end of naïve period), and/or (ii) a predation shift by wolves, which targeted their predation on red deer instead of roe deer (Appendix 3) with increasing pack size.

## Acknowledgements

We sincerely thank the successive Presidents of the “Fédération Départementale des Chasseurs de la Drôme” (Alain Golin, Alain Hurtevent, Rémi Gandy), the Director of the “Fédération Départementale des Chasseurs de la Drôme” (Denis Rix) and all board members of the “Fédération Départementale des Chasseurs de la Drôme” for providing all means (technical, funding, staff) to carry out this research. We also thank the ONF technicians of the Vercors and Lente-Léoncel for their invaluable help with data collection and our fruitful discussions, particularly Jean-Pierre Lacour, Eric Rousset, Yves Pesenti, Gaël Gauthier, Franck Millat Carus, Philippe Joanin and Thierry Rozand. We are indebted to all the hunters, and particularly to all the Presidents of the ACCA, without whom this study would not have been possible. We thank Simon Chamaillé-Jammes for fruitful discussions about the interpretation of the results of this study, and Eric Marboutin and Maryline Pellerin for comments on the manuscript. Finally, we thank Jean-Pierre Fermond (“Bubu”), Jérôme Guilloud and Christophe Randon who have supported this study since the beginning, provided guidance throughout and commented earlier versions of this draft.

**Author contributions**
MR conceived and designed the study. MR, JM and TC coordinated the fieldwork. CB and MV analyzed the data. MR, CB, CT, JMG and MV interpreted the results. CB and MV wrote the initial manuscript. All authors revised the manuscript.

